# Whole-genome sequencing analysis of anthropometric traits in 672,976 individuals reveals convergence between rare and common genetic associations

**DOI:** 10.1101/2025.02.24.639925

**Authors:** Gareth Hawkes, Harrison I. W. Wright, Robin N. Beaumont, Kartik Chundru, Aimee Hanson, Leigh Jackson, Anna Murray, Kashyap Patel, Timothy M. Frayling, Caroline F. Wright, Andrew R. Wood, Michael N. Weedon

## Abstract

Genetic association studies have mostly focussed on common variants from genotyping arrays or rare protein-coding variants from exome sequencing. Here, we used whole-genome sequence (WGS) data in 672,976 individuals of diverse ancestry to evaluate the contribution and architecture of rare non-coding variants to three commonly studied anthropometric traits: height, body mass index (BMI) and waist-hip ratio adjusted for BMI (WHRadjBMI). Analysing 447,461 individuals in UK Biobank for discovery and 225,515 individuals in All of Us for replication, we identified 90 novel rare and low-frequency single variant associations. This includes two independent rare variants upstream of *IGF2BP2* that both substantially reduce WHRadjBMI, but have distinct effects on other adiposity traits. We identified 135 coding variant aggregates, several of which were missed by exome sequencing studies. For example, *UBR3* protein-truncating variants were associated with a 2.7kg/m2 increase in BMI. We additionally identified 51 non-coding variant aggregate associations, including in the 5’UTR of *FGF18* (a highly constrained gene with no previously reported coding associations) associated with up to 6cm effects on height. We show that 97% of rare variant associations occur near GWAS loci demonstrating convergence of rare and common variant associations. Finally, we show that ultra rare variants (MAF<0.01%) explain a small fraction of heritability (<10%) compared to common variants for these traits, that heritability is largely shared across ancestries, and that this heritability is concentrated at or near common variant loci. Our work demonstrates the importance of large-scale WGS for fully understanding the genetic architecture of complex traits.

## Introduction

Human genetic association studies have mostly focussed on common variants from genome-wide association studies (GWAS), or rare protein-coding variants from exome sequencing studies. Despite the successes of these studies, a substantial amount of heritability remains unaccounted for^1^. For example, a GWAS of 5.4 million individuals identified 12,111 common variant associations for adult height^2^, which accounted for ∼40% of the variance among individuals of European genetic ancestry. However, the total heritability for height is estimated to be ∼80%^3^, indicating that many additional genetic factors remain to be discovered.

Rare non-coding variants may explain a substantial amount of the remaining heritability, but their contribution is largely unexplored. The causal variants and effector genes at most GWAS loci have not been identified using imputed and exome sequencing data because of small effect sizes, extensive linkage disequilibrium (LD) and because most associations are non-coding. Larger effect sizes and limited LD mean that rare variants have substantial advantages over common variants in identifying causal variants and genes. Rare non-coding associations will therefore allow us to identify key regulatory elements for genes important in human biology and disease^4^. Identifying rare non-coding associations may also substantially aid the identification of the causal gene at GWAS loci.

There are many examples of genes where rare coding variants cause monogenic disease while common, usually non-coding, variants subtly predispose to a related trait. For example, loss of function variants in *MC4R* cause severe obesity^5^, whereas distal non-coding regulatory variants are associated with a small increase in BMI^6^. A spectrum of associations from common low-effect to rare large-effect variants at a locus (an ‘allelic series’) can substantially aid in the identification of the effector gene at a locus^7^. It also provides the opportunity to assess the effect of differential levels of disruption on a gene which can provide key insights into biological mechanism and can be important in drug development^8^. How often rare and common variant associations converge on the same genes^9^ and regulatory elements is therefore a key unanswered question.

WGS-association analyses in 100,000’s of individual’s is now becoming possible. For example, the UK Biobank (UKB) has released WGS data from 500,000 individuals^10^ and, at the date of writing, the All of Us study^11^ (AoU) has WGS data available on nearly 250,000 individuals. We previously used an interim release of WGS data on 200,003 individuals from UKB to develop a framework for WGS-based association testing^12^. We discovered and replicated rare variant associations missed by single variant-based GWAS and exome sequencing studies. For example, we identified rare non-coding aggregate-based associations for height proximal to *HMGA1* and *miRNA-497*.

Here, we substantially extend our previous study by analysing 447,461 individuals in UKB for three traits that have been the central focus of previous large-scale GWAS meta-analyses: adult height, body-mass-index (BMI), and waist-hip-ratio adjusted for BMI (WHRadjBMI). We replicate our findings in 225,515 individuals of diverse genetic ancestry from the AoU study. We identify hundreds of novel rare non-coding and rare protein-coding variant associations; identify many rare non-coding variant associations that can only be identified from WGS analysis; demonstrate the power of WGS analyses to fine-map common variant associations; and show that, for these traits, most rare variant heritability co-localises with GWAS loci.

## ONLINE METHODS

We performed primary discovery association analyses for three key anthropometric traits: height, body mass index (BMI), and waist-hip-ratio adjusted for BMI (WHRadjBMI) using annotated variants from WGS data on up to 447,461 individuals of inferred European (EUR) genetic ancestry from the UKB, a population cohort from the United Kingdom. As our interest was in novel rare variants, we did not consider the two second-largest ancestries within UKB due to their small sample size (South Asian; UKB-SAS & African; UKB-AFR; N<10,000). We replicate our results in the diverse AoU Cohort, based on individuals of inferred European (AoU-EUR; N=128,566), African (AoU-AFR; N = 54,940) and admixed-American (AoU-AMR; N = 42,009).

We performed both single variant (minor allele count (MAC) ≥ 5) and genomic aggregate association tests (minor allele frequency (MAF) < 0.1%) using REGENIE^13^. Phenotypes were adjusted for age, age squared, sex, recruitment centre, WGS centre and 40 genetic principal components at runtime (**Methods**), and identified independent variant associations using a modified version of GCTA-CoJo^14^. We annotated all genetic variants using Ensembl’s Variant Effect Predictor (VEP)^15^ (**Methods)** and used the output to categorise variants as gene-centric (e.g., coding, predicted intronic splicing, intronic unspecified, proximal-regulatory) and intergenic-regulatory (e.g., Ensembl regulatory regions, non-coding RNA, intergenic unspecified) for aggregate-based association testing. Additionally, we performed aggregate testing on all non-coding variants in overlapping (1kbp overlap) 2kbp sliding windows. We also sub-categorised variants within a subset of aggregate units by measures of constraint (JARVIS^16^), conservation (GERP^17^) and/or predicted deleteriousness (CADD^18^).

## RESULTS

### 160 rare and low frequency variants associated with anthropometric traits in UKB after adjusting for known variants

We identified a total of 2,690 statistically independent single-variant associations in our UKB-based analysis for height, BMI and WHRadjBMI reaching P<3×10^−10^ (**Supplementary Figure 1**). Of the 2,690 genetic associations, 91 were rare (MAF < 0.1%) and 114 were low-frequency (0.1% ≤ MAF <1%) (**Supplementary Table 1**). After adjustment for variants previously reported to be associated with the respective trait by the GIANT consortium^2,19–23^, 215 (8%) single-variant associations remained significant, of which 160 (74.4%; 137 for height, 13 for BMI and 10 for WHRadjBMI) were low-frequency or rare (MAF < 1%; **Supplementary Table 2**).

### Single-variant associations replicate in AoU with effect sizes consistent across ancestries

We attempted to replicate our results using 225,515 individuals from the AoU study (N EUR=128,566; N AFR=54,940; N AMR=42,009). Of the 215 (common, low-frequency and rare) single-variant associations in UKB, 172 were available for analysis with minor allele counts (MAC)≥5 in one or more of the three broad genetic-ancestry groups in AoU (42 with UKB-MAF < 0.1%; 77 with 0.1% ≤ UKB-MAF < 1%; 53 with UKB-MAF ≥ 1%; **Supplementary Table 3**). Of the 119 novel rare and low-frequency variant associations which we could put forward for replication, 90 (75.6%) showed nominal evidence of replication (*P* < 0.05), and 45 (37.8%) showed strong evidence of replication at *P* < 0.05/119. Of the 53 common variants with UKB-MAF > 1% put forward for replication, we observed 48 (90.6%) associations with nominal evidence of replication (P < 0.05), and 35 (66.0%) with Bonferroni-corrected evidence of replication (P < 0.05/53). No variants with UKB-MAC ≤ 10 (7/172; 4.07%) showed even nominal evidence of replication. However, we observed strong directional consistency in effect sizes for all single variants reaching P < 0.05 across ancestries (**Figure 1**: N directionally consistent = 137/138; 99.3%; Binomial *P* = 7.53 × 10^−177^). Fifteen percent (33/215) of these variants would not have been detected by conventional GWAS and imputation-based approached because they were not present in the Haplotype Reference Consortium + UK10K, TOPMed, or Genomics England imputed UKB datasets available in UK Biobank RAP. Of the remaining 82 (85%) variants, the imputation quality across the imputation panels was variable (**Supplementary Table 4**).

**Figure 1:**
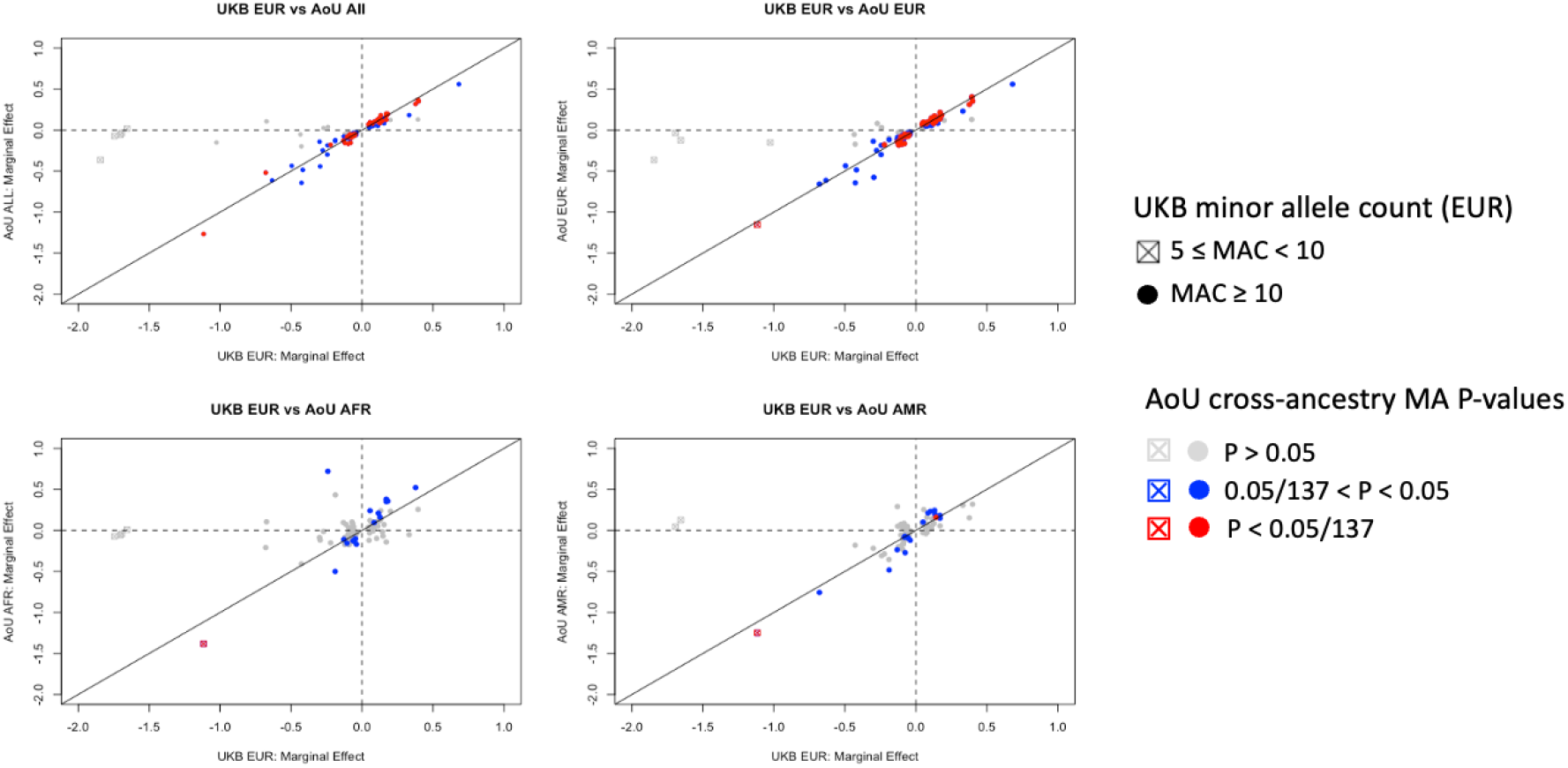
Replication of UK Biobank single variant associations in All of Us. Comparison of single variant effect estimates from discovery (UK Biobank European; UKB-EUR) to replication analyses (All of Us European, Admixed-American and African; AOU-EUR, AOU-AMR and AOU-AFR respectively). Points are coloured according to their p-value. Circles represent cases where UKB-EUR minor-allele-count (MAC)>20, and squares represent variants where UKB-MAC≥5. Only variants with MAC≥5 in at least one AOU ancestry were put forward for replication.

#### A single rare variant in the seed region of MIRNA497 affects height by 4cm

We identified a rare insertion (GRCh38:17:7017939:C:CT; MAF = 0.014%) in *MIRNA497 (P* = 6.53 × 10^−16^), with a per allele effect of 3.79cm [95% CI 2.87, 4.71cm] on height in UKB. This association was replicated in AoU (*P* = 1.38 × 10^−27^). This variant partially explains an aggregate-based association we previously discovered of highly conserved variants in and near miRNA host-gene *MIRNA497HG* and its by-products *MIR195* and *MIR497* ^12^. However, this insertion is statistically independent of an aggregate-based association of highly-conserved (GERP>2) variants overlapping the promoter region of host-gene *MIR497HG* (*P* = 7.17 × 10^−12^), i.e. the remainder of the signal. Our results suggest that regulation of the miRNA497 product impacts height independently of the coding consequences acting via the rare insertion.

#### Two independent rare variants upstream of IGF2BP2 associate with WHRadjBMI

We identified two independently-associated rare single variants upstream of *IGF2BP2* associated with WHRadjBMI: GRCh38:3:185826396:AG:A (beta = -0.17 SD [-0.22, -0.12], MAF = 1.59 × 10^−3^, *P* = 2.12 × 10^−11^, replication *P* = 2.24 × 10^−2^) and GRCh38:3:185847637:G:A (beta = -0.122SD [-0.158, - 0.086], MAF = 3.10 × 10^−3^, *P* = 2.99 × 10^−11^, replication *P* = 4.19 × 10^−4^). Although both variants associate strongly with WHRadjBMI, they have heterogenous effects on other phenotypes. For example, the former occurs in a lncRNA (TCONS_00006340) previously linked to fasting glucose^24^ and is associated with favourable adiposity and height, whilst the latter is not (**Figure 2; Supplementary Table 5**), suggesting different regulatory effects on *IGF2BP2*.

**Figure 2:**
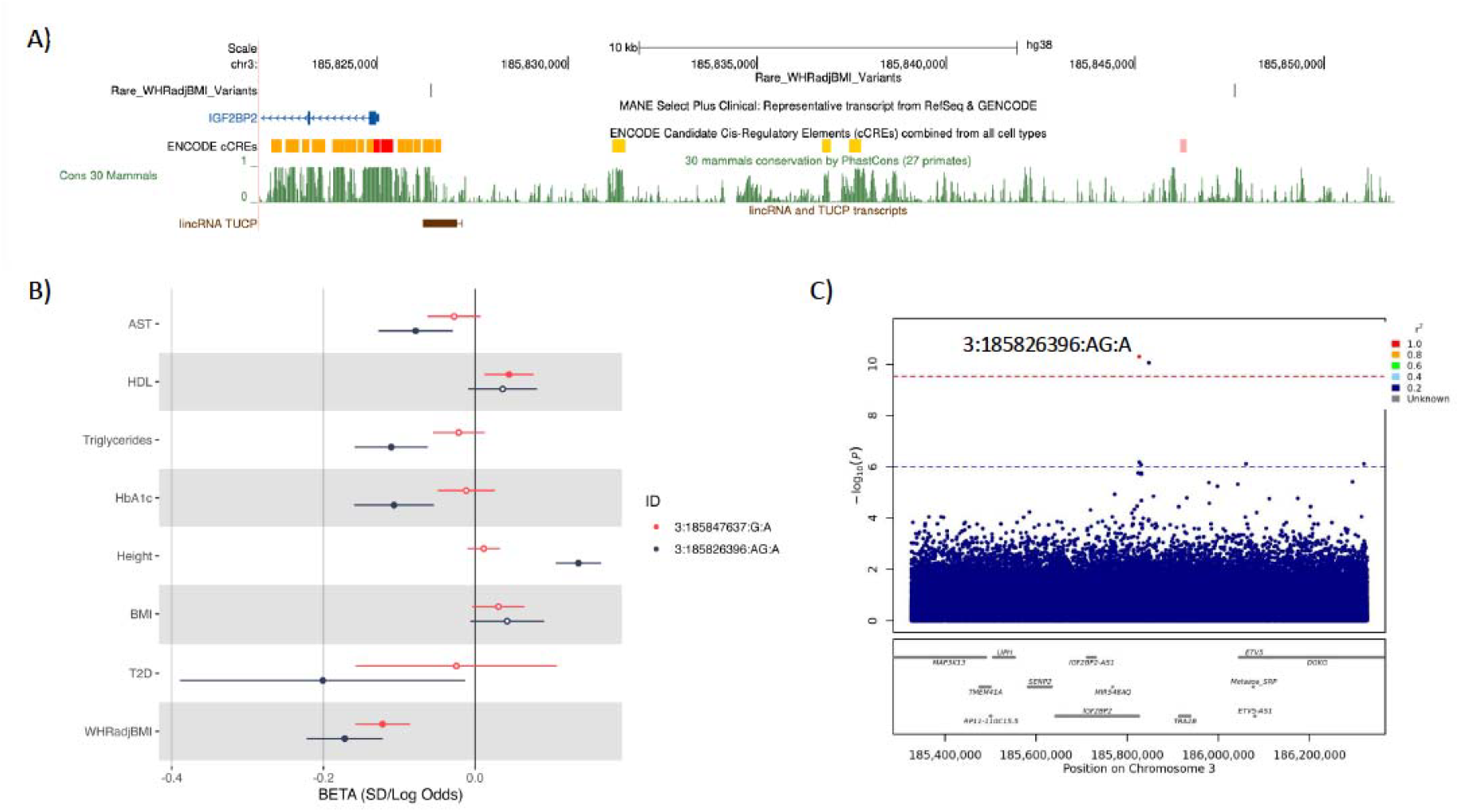
Summary of genetic associations at the *IGF2BP2* locus with WHRadjBMI. A) UCSC genome browser plot showing the location of the two rare variants relative to *IGF2BP2*, a measure of conservation (Cons 30 Mammals) and the overlapping lincRNA. B) Forest plot showing the effect size of 3:185847637:G:A and 3:185826396:AG:A against metabolic-related phenotypes. C) LocusZoom plot, including common variants, centred on the lincRNA-overlapping variant (purple diamond = 3:185826396:AG:A).

### WGS identifies protein-coding aggregate-variant associations missed by exome sequencing

We identified 135 independent rare coding variant (MAF < 0.1%) aggregate associations (**Supplementary Table 6**), of which 125 associations (height: 114; BMI: 8; WHRadjBMI: 3) remained significant after adjustment for variants previously reported by the GIANT consortium as associated with the respective trait **(Supplementary Table 7**). The set of genic aggregate-based associations included 73 predicted loss-of-function (58.4%), 49 missense (39.2%), 2 splice region (1.60%) and 1 synonymous (0.80%). We put forwards 118 of the coding aggregate-based associations for replication in AoU where there were sufficient numbers of carriers, defined as at least five minor-allele-carriers in any of the three largest ancestral groups. Of the 118 associations, 91 showed nominal (*P* < 0.05) evidence of replication, and 45 passed a Bonferroni-corrected P-value threshold of *P* < 4.24×10^−4^ (**Supplementary Table 8**). The one synonymous aggregate association (in *IRS4*, associated with height) did not replicate, suggesting a likely false-positive result.

Notable protein-coding aggregate-based associations, which were not significantly associated in existing exome-based studies, include predicted loss-of-function variants in *PCSK5 (P*=1.19×10^−19^, *P* replication=3.76×10^−3^), *AMOTL2* (*P=2*.21×10^−9^, *P* replication=0.02), *BCL9 (P*=5.69×10^−9^, *P* replication=0.02), *MAZ* (*P*=1.24×10^−10^, *P* replication=0.01), and *TRPC4AP* (*P=3*.41×10^−12^, *P* replication=0.04) associated with height. We also observed novel coding aggregate-based associations for predicted highly-deleterious missense variants (CADD>25) in *AXL* (*P*=5.39×10^−11^, *P* replication=5.72×10^−4^), *LRRC58* (*P*=1.21×10^−15^, *P* replication=0.01), and *SMAD3* (*P*=3.87×10^−9^, *P* replication=0.01).

We further identified an association between predicted loss-of-function variants in *UBR3* and BMI (*P*=1.21×10^−9^, *P* replication=3.12×10^−3^) that was not reported by recent UKB exome sequencing association studies. Although the *UBR3*-BMI association was strongest in a *SKAT* framework, which accounts for bi-directional effects, we also observed a genome-wide significant burden (additive) effect (beta = 2.64kg/m2 [3.51, 1.78], *P* = 2.20×10^−9^).

### We identified 51 non-coding aggregate associations across anthropometric traits, particularly in 5’UTRs

We identified 53 independent rare-variant aggregate non-coding associations (**Supplementary Table 9**), 51 of which remained significant after adjusting for previously reported loci by the GIANT consortium **(Supplementary Table 10**): 26 for height and 25 for BMI. In total, we identified six 5’ untranslated regions (5’UTR; 11.8%) and four upstream (7.84%) associations, as compared to two 3’UTR (3.92%) and three downstream (5.88%). We additionally identified three ENSEMBL regulatory-region aggregate associations, and four regions annotated as RNA. We observed an enrichment of 5’ UTR associations compared to the frequency of all 5’UTR aggregates tested (background = 2.78%; Fisher’s Exact OR = 4.66, *P* = 2.85 × 10^−3^). Of the 51 associations, we were able to test 50 for evidence of replication in All of Us (**Supplementary Table 11**): 6 showed nominal evidence of replication, and 3 replicated at a Bonferroni-corrected threshold (*P <* 0.05/50).

### Rare 5’UTR variants in FGF18 substantially influence height

We identified a novel non-coding aggregate-based association between 5’UTR variants in *FGF18* and height in UKB (*P* = 2.93 × 10^−12^), with evidence of replication in AoU *(P* = 1.16 × 10^−4^). In our analysis, coding variants of any consequence in *FGF18* were not associated with height (*P* ≥ 2.12 × 10^−3^), and associations were also absent in both Genebass^25^ and the Astrazeneca PheWAS portal^26^.

Further analyses suggested that this aggregate-based association was not driven by any single variant (min single variant association *P* ≥ 6.34 × 10^−8^). Within the aggregate, we observed eight variants with MAC ≥ 5, presenting with effect sizes ranging from 6.71cm to –2.94cm (*P*<0.05; **Supplementary Table 12; Figure 3**). Four of those eight variants were located at a single multi-allelic position (GRCh38:5:171420095), overlapping an enhancer. Previous GWAS efforts have highlighted seven statistically independent common (MAF >1%) variants in a cross-ancestry meta-analysis^2^ with *FGF18* as their closest gene, and effect sizes ranging from –0.26cm to 0.13cm. Together, these results provide a clear example of using rare-variant aggregate testing for the purposes of fine-mapping previously reported GWAS signals.

**Figure 3:**
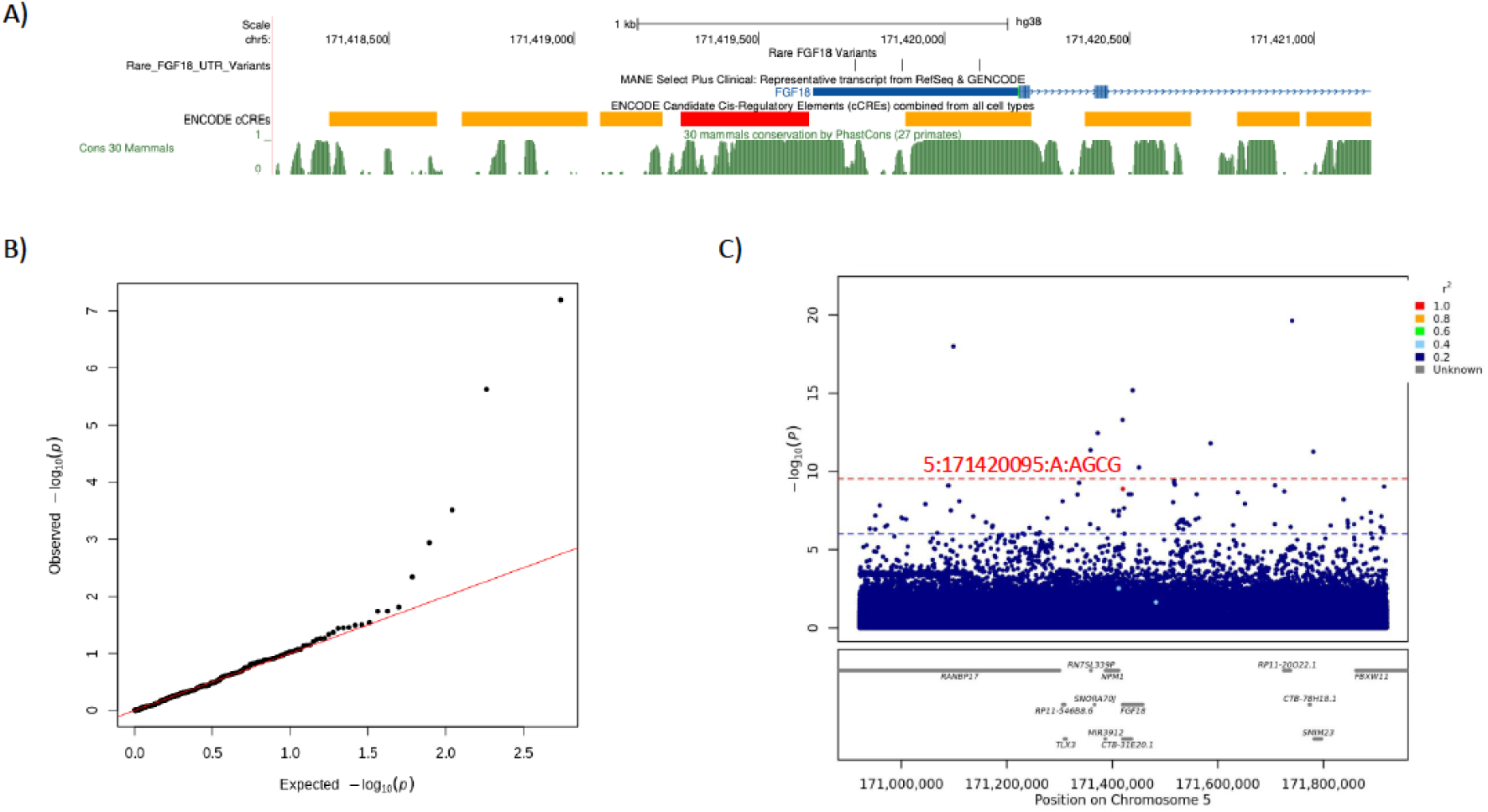
Summary of association between rare variants in the 5’UTR of *FGF18* and height. A) UCSC genome browser plot showing the *FGF18* region highlighting the position of the most strongly associated single variants in the aggregate, ENCODE cCREs and Conservation (Cons 30 Mammals). B) QQ plot of variants contributing to the *FGF18* proximal association. C) LocusZoom plot, adjusted for common (>1% MAF) lead variants, centred on the most-strongly associated variant in the aggregate (5:171420095:A:AGCG).

### Rare single variant and aggregate-variant associations with anthropometric traits usually occur near GWAS loci

We assessed the proximity of rare single variant and aggregate-variant height associations with common variants identified from GWAS. We focussed on height firstly because common variant heritability has been saturated for European individuals as presented in Yengo et. al 2022^2^, and secondly due to the large number of rare single variant and aggregate-variant associations. We assessed proximity to 12,111 common variant associations that were reported to saturate common SNP-based heritability in Europeans and span ∼21% of the genome based on 70kb windows centred on sentinel SNPs (±35kb)^2^. Of the 75 variant associations with UKB-EUR MAF <1% that remained statistically significant in our European-based UK Biobank analysis and with P<0.05 in AoU-EUR, 64 (85%) and 73 (97%) resided within 35kb and 100kb (∼44% of the genome), respectively (**Figure 4**). These observations were similar for 35 common variants that remained statistically significant with 32 (91%) and 33 (94%) residing within 35kb and 100kb of the 12,111 SNPs, respectively (**Supplementary Table 13**). Of the 73-coding aggregate-based associations that remained statistically significant in our UKB-EUR-based analysis and with P<0.05 in AoU-EUR, 61 (84%) and 70 (96%) resided within 35kb and 100kb of the 12,111 SNPs, respectively. Of the seven autosomal non-coding aggregate-based associations that remained and with P<0.05 in AoU-EUR, all were located within 35kb of a previously reported common variant (**Supplementary Table 14**).

**Figure 4:**
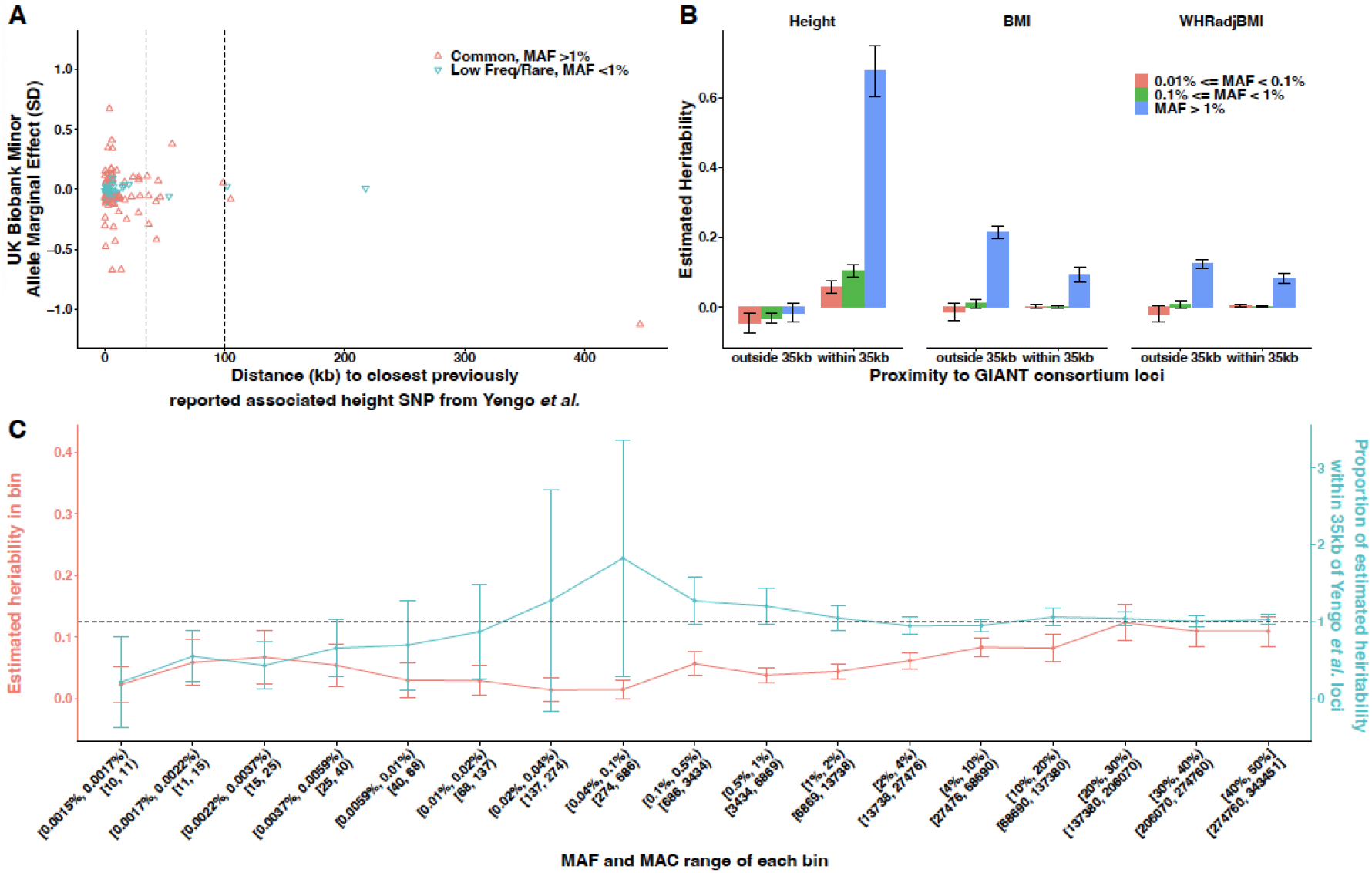
Co-localisation and heritability analyses of anthropometric traits. A) Co-localisation of rare (MAF<1%) and common (MAF>1%) independent single variants associated with height in our analysis, relative to those presented in the common-variant heritability saturation *Yengo et. al 2022*. **B)** Estimates of heritability for three variant-frequency bins within and outside of 35kbp of common variants reported by the most recent GIANT analysis of height, BMI and WHRadjBMI. **C)** Semi-continuous estimation of heritability for height across 17 variant frequency bins, in contrast to the proportion of heritability for each bin within 35kbp of a *Yengo et. al 2022* variant.

### Rare single variant and aggregate-variant associations also converge across ancestries

Next, we assessed the proximity of genetic associations for height in other broad genetic ancestries, using 54,940 and 42,009 individuals of African (primarily African-American) (AFR) and admixed-American (AMR) genetic ancestry, respectively, in AoU. In our AFR-based analysis, we identified 52 variants associated with height, of which 45 (87%) reside within 35kb of a previously reported common variant, including 3/5 associated variants with AoU-AFR MAF <1%. We also observed 49/52 (94%) associated variants reside within 100kb, including 4/5 (80%) with AoU-AFR MAF <1% (**Supplementary Table 15**). A similar pattern of physical colocalization was observed in our AMR-based analysis. Of the 35 variants identified as associated with height, 30 variants (86%) reside within 35kb of a previously reported common variant, including 2 of 3 (67%) with AoU-AMR MAF <1%. In addition, 34/35 (97%) reside within 100kb of a previously reported common variant, including all associated variants with AoU-AMR MAF <1% (**Supplementary Table 15**).

### Heritability explained by rare variants is enriched around common signals of association

To determine whether rare and common genetic signals of association are likely to converge independently of the genetic associations we had power to detect, we performed partitioned heritability analyses using RHEmc (**Methods**). Using a maximally unrelated subset (common-variant GRM off-diagonal < 0.05) of the UKB-EUR WGS-based analysis (N = 343,451), we observed a total h2 estimate of 0.744, 0.309, and 0.202 (se = 0.036, 0.013, 0.016), of which 11.2%, 0.1%, and –3.4% (h2=0.084, 0.0004, -0.007 (se = 0.021, 0.014, 0.013)) is explained by low-frequency and rare variants (UKB-EUR MAF < 1%), for height, BMI, and WHRadjBMI, respectively. Of the heritability explained by the low-frequency and rare variants, 100% can be attributed to the variants residing within 35kb of a previously reported common variant associated with height^2^ (**Figure 4, Supplementary Table 16**). This value was lower for BMI and WHR adjusted for BMI (31% and 44% respectively), which may reflect the potential exclusion of 35kb windows around common GWAS signals that are yet to be detected through larger-scale GWAS meta-analyses for these traits.

Finally, we performed a secondary heritability analysis for height to additionally stratify by allele frequencies of variants in the WGS data. We used 17 minor-allele frequency bins, down to and including variants with a minor-allele count of 10, and three LD bins per frequency bin (**Figure 4**; **Methods**). The estimate for the proportion of heritability within 35kb of a previously reported height GWAS SNP was > 0.8 for 12/17 bins, and >0.99 for 9/17 bins. However, the total heritability estimate summed across all 17 frequency bins was inflated (h2 = 1.005, SE = 0.0535), with inflation observed across rare-frequency bins (potentially indicative of residual population stratification). We quantified the inflation in the ultra-rare (MAF<0.0001) variant bins using two orthogonal approaches. First, we fitted a logarithmic model on the cumulative heritability of the bins used in the primary analysis (MAF>1e-4; **Supplementary Figure 2**) which gave an estimate of h2 = 0.8646, which would imply an inflation of 0.1403 from the ultra-rare bins. Secondly, assuming all heritability outside of the 35kb regions is inflated, we adjusted the estimate within the 35kb regions for the ultra-rare bins by the estimate outside of the 35kb regions (**Supplementary Table 17**) adjusted for the total bases covered by the regions (21% within 35kb vs 79% outside). We observed h2=0.8648, implying a total inflation of 0.1401. Both methods estimate a heritability of the ultra-rare variants of h2=0.094, which would likely represent an upper-bound for the true value.

Together, these results indicate that rare variants associations are most likely to converge with common genetic signals of association for anthropometric traits.

## Discussion

Here, we use large-scale WGS association analyses to discover hundreds of new rare variant associations for anthropometric traits. We identify rare single variants that have been missed by previous GWAS studies based on imputation. Using rare variant aggregates, we then identify gene-phenotype associations missed by exome sequencing studies and provide clear examples of non-coding aggregate-variant associations for genes that have no coding associations. Finally, our results suggest that rare single variant, coding and non-coding variant aggregates converge on the same genes.

We demonstrate that the majority of previously unreported rare variants associated with height in an analysis of WGS data co-localise with common variants reported by the most recent meta-analysis of imputed data, which reported saturation of the common-variant heritability among individuals of European ancestry. We further show that this result extends to other genetic ancestries, which did not experience a similar heritability saturation. In previous work, Wu et al.^27^ inferred through simulations that co-localisation patterns of 35kb to causal variant for >80% of signals only applied when simulating within frequency bins, i.e. they observed common variants to reside close to common causal variants and rare signals to reside close to rare causal variants. Our work demonstrates that both common and rare variant associations converge on the same genomic regions and genes. This finding is important because it demonstrates the underlying biological pathways are likely to be similar and we are therefore assessing allelic series of effects on genes. Our results also suggest these pathways are consistent across genetic ancestries. Finally, restricting the hypothesis-space for an analysis to the region around known GWAS loci could substantially reduce both multiple-testing thresholds, and the environmental impact of computing resources required to analyse these large-scale genetic data^28^.

Our work provides another clear example of a non-coding association near a gene that lack coding associations, specifically, the association of rare 5’UTR variants in *FGF18* with height. This is a small, constrained gene (pLI=0.98; missense Z=2.44)^29^ with only three protein truncating variants in UK Biobank, none of which occur in the last exon where they would be expected to escape nonsense-mediated decay. This result is similar to a previous finding in *HMGA1*, where we previously showed that rare regulatory variants affect height by up to 5 cm, but there is no protein coding association for the gene. Coding associations in both these genes may be undetectable, either due to lack of power or because haploinsufficiency (or other coding effects) is incompatible with life, whilst more subtle effects on expression are compatible with life and result in a detectable effect on height. Both *FGF18* and *HMGA1* have common variant GWAS signals, and our work demonstrates that these are the likely causal genes.

We further demonstrate that aggregate testing is a powerful way to identify novel associations with rare coding and non-coding variants. We previously identified *MIRNA497* as being associated with height using an aggregate-based association in 200,000 WGS in UK Biobank^12^. With increased statistical power we now show a single rare deletion in the seed region of *MIRNA497* is independently associated with height and is a likely causal variant at the locus. Our results now suggest that some variants influence the expression of the *MIRNA497* and others independently influence height by affecting its ability to bind targets. There is substantial orthogonal evidence for an important role of *MIRNA497* in human growth^30–32^.

We identify several novel anthropometric trait gene associations, missed by existing exome sequencing studies. We found a substantial number of novel coding aggregate associations for height. Most of these missed associations were because of low coverage in the exome sequencing compared to genome sequencing. For example, one of the most strongly associated genes is *LCORL*, a GWAS locus for height. In GeneBass and Astrazeneca PheWAS portal *LCORL* PTVs are not associated with height (P > 0.01), but in our WGS analysis it associates with P < 1×10^−60^ in UK Biobank and P < 1×10^−17^ in All of Us. This is, at least partly, explained by poor coverage of some of the exons in *LCORL* from exome sequencing. Other explanations include, e.g., poor array-based genotyping/imputation of rare variants, poor quality exome sequencing in GC-rich exons, lack of methods for meaningful aggregate testing, etc.

There are a number of limitations to our study. We did not explore physical colocalization of common and rare genetic associations for BMI and WHRadjBMI for two primary reasons. First, in contrast to height, we identified relatively few rare genetic associations for BMI and WHRadjBMI. Second, more common SNP-based heritability remains to be explained in all broad genetic ancestry groups.

Consequently, rare genetic associations may reside close to common variant associations that are yet to be detected by larger GWAS efforts. Furthermore, both UKB and AoU are population cohorts that are affected by healthy volunteer recruitment bias and may therefore be depleted of individuals with more extreme phenotypes. Finally, WGS data were generated using short-read sequencing, so information on structural variants and haplotypes was missing. Future studies of more diverse ancestries and using long-read sequencing technologies are likely to add further insights to the field. We additionally did not consider individuals of non-EUR UKB participants due to sample size constraints.

In summary, we have shown that WGS enables the discovery of novel associations with rare variants not found by other technologies. We have found novel coding associations for several anthropometric traits in genes with poor exome-sequencing coverage, and non-coding associations near genes for which no coding association could be identified. Furthermore, we have shown that common, rare and aggregate associations for height converge on the same loci, suggesting shared underlying biology.

## Acknowledgements, Funding, Author Contributions

RNB, GH and MNW are supported by Medical Research Council grant MR/Y003748/1. ARW is supported by the Academy of Medical Sciences / the Wellcome Trust / the Government Department of Business, Energy and Industrial Strategy / the British Heart Foundation / Diabetes UK Springboard Award [SBF006\1134]. TMF is supported by MRC awards MR/WO14548/1 and MR/T002239/1. The research utilised data from the UK Biobank resource carried out under UK Biobank application number 103356. UK Biobank protocols were approved by the National Research Ethics Service Committee. The authors would like to acknowledge the use of the University of Exeter High-Performance Computing (HPC) facility in carrying out this work, funded by an MRC Clinical Research Infrastructure award (MRC Grant: MR/M008924/1). This study was supported by the National Institute for Health and Care Research Exeter Biomedical Research Centre. The views expressed are those of the authors and not necessarily those of the NIHR or the Department of Health and Social Care. We gratefully acknowledge All of Us participants for their contributions, without whom this research would not have been possible. We also thank the National Institutes of Health’s All of Us Research Program for making available the participant data examined in this study

## Data Availability

Data cannot be shared publicly because of data availability and data return policies of the UK Biobank and All of Us. Data are available from the UK Biobank for researchers who meet the criteria for access to datasets to UK Biobank (http://www.ukbiobank.ac.uk) and All of Us (https://allofus.nih.gov/).

## Methods

### UK Biobank and All of Us Whole Genome Sequencing

The whole genome sequencing performed for UKB had an average coverage of 32.5X using Illumina NovaSeq 6000 sequencing machines^10^. The genome build used for sequencing was GRCh38: single variant nucleotide polymorphisms and short ‘indels’ were jointly called using DRAGEN 3.7.8.

We set any UKB-WGS genotype calls to missing if either the sum(LAD)<8 (local allele depth; LAD) per sample-genotype or GQ<10 (genotype quality; GQ) for each of the 154,430 pVCFs provided by UKB using bcftools^33^. After these additional quality control steps, the transmission rate of singletons, which should theoretically be exactly 0.5 (assuming the majority of variants are not under strong negative selection), was 0.497, as compared to 0.456 as originally provided by UKB.

The whole genome sequencing performed for AoU had average coverage ≥30X using Illumina NovaSeq 6000 sequencing machines. The genome build used for sequencing was GRCh38: single variant nucleotide polymorphisms and short ‘indels’ were jointly called using DRAGEN 3.4.12.

We exported the All of US WGS data from the provided VDS format to VCF format using HAIL v0.2.126. We subsequently set genotype calls to missing if ‘FILTER!=PASS’ (AoU site level filtering; FILTER), and, as with the UKB WGS data, set REF/ALT and ALT/ALT genotype calls to missing if either the sum(LAD)<8 or GQ<10 per sample-genotype, using bcftools^33^. We were unable to filter homozygote reference calls in the AOU WGS data because those LAD values were not provided, and those GQ values were not provided at the base-pair level.

For both All of Us and UK Biobank, we subsequently dropped any variant with a missingness greater than 10%. Indels were normalised and left-aligned using bcftools based on a 1000G b38 reference available at https://ftp.1000genomes.ebi.ac.uk/vol1/ftp/technical/reference/GRCh38_reference_genome/ (accessed 30/03/2024). Finally, a multi-allele splitting procedure was applied, and each variant was assigned a unique ID (CHR:BP:REF:ALT) before merging all VCFs per chromosome. Each merged pVCF was then converted to plink^34^ (v2.0) p(gen/var/sam) format.

### Genetic Variant Annotation

We annotated all genetic variants using Ensembl Variant Effect Predictor (VEP)^15^, LOFTEE^35^ and UTRannotator^36^. Where possible, we assigned each variant to one of three *classifications*: coding, proximal-regulatory or intergenic-regulatory. A variant was classified as coding if it had a predicted impact on the coding sequence of **any** transcript; proximal-regulatory if the variant lay within a 5kbp window of the UTRs of a transcript, and was not already a coding variant in any transcript, and finally regulatory if was not coding, but we did not assume proximality to coding transcript (see below). We additionally tested variants in sliding windows of size 2000 base pairs, regardless of the number of variants in each window, with coding and proximal-regulatory variants excluded to minimise hypothesis-testing overlap.

We then assigned each variant to groupings, which we refer to as *masks*, according to their predicted consequence and location. We used five published variant scores to group variants by consequence:

1. **Genomic Evolutionary Rate Profiling (GERP)** The GERP score is a measure of conservation at the variant level^17^. We classified a variant as highly conserved if it had a GERP score >2.
2. **phastCon score** phastCon is a window-based measure of conservation across species^37^: either strictly mammalian (phastCon 30), or for all species (phast_100). We tested non-coding genome windows, i.e. excluding any window containing an exon, that had a phastCon score in the 99^th^ percentile.
3. **Constraint Score** Constraint was calculated in windows of size 1kbp^38^ based on the local mutability and observed mutation rate of each window. We tested windows with a constraint z-score greater than or equal to four.
4. **SpliceAI score** The SpliceAI score^39^ is a measure of how well predicted each variant within a pre-mRNA region is of being a splice donor/acceptor, or neither. A variant was classified as a splice site with high confidence if it had an AI>50.
5. **Combined Annotation Dependent Deletion score (CADD)** The CADD score^18^ predicts how deleterious a variant is likely to be. We applied the CADD score only to coding variants and considered loss-of-function variants only if tagged as high confidence by VEP. Missense variants with CADD>25 were segregated for testing in a separate mask.
6. ***JARVIS* Score** The JARVIS score^16^ was derived to better prioritise non-coding genetic variation for association study, based on a machine learning model derived from measures of constraint.

Each genome mask consisted of a number of variants with different *consequences*, based on their location, one of the above scores and/or predicted coding consequences. For example, for a variant to be classified as missense CADD>25, it must change a codon of an exon of a gene transcript and be predicted to be highly deleterious.

In Table 1 we present the full list of consequences assigned to each mask and classification.

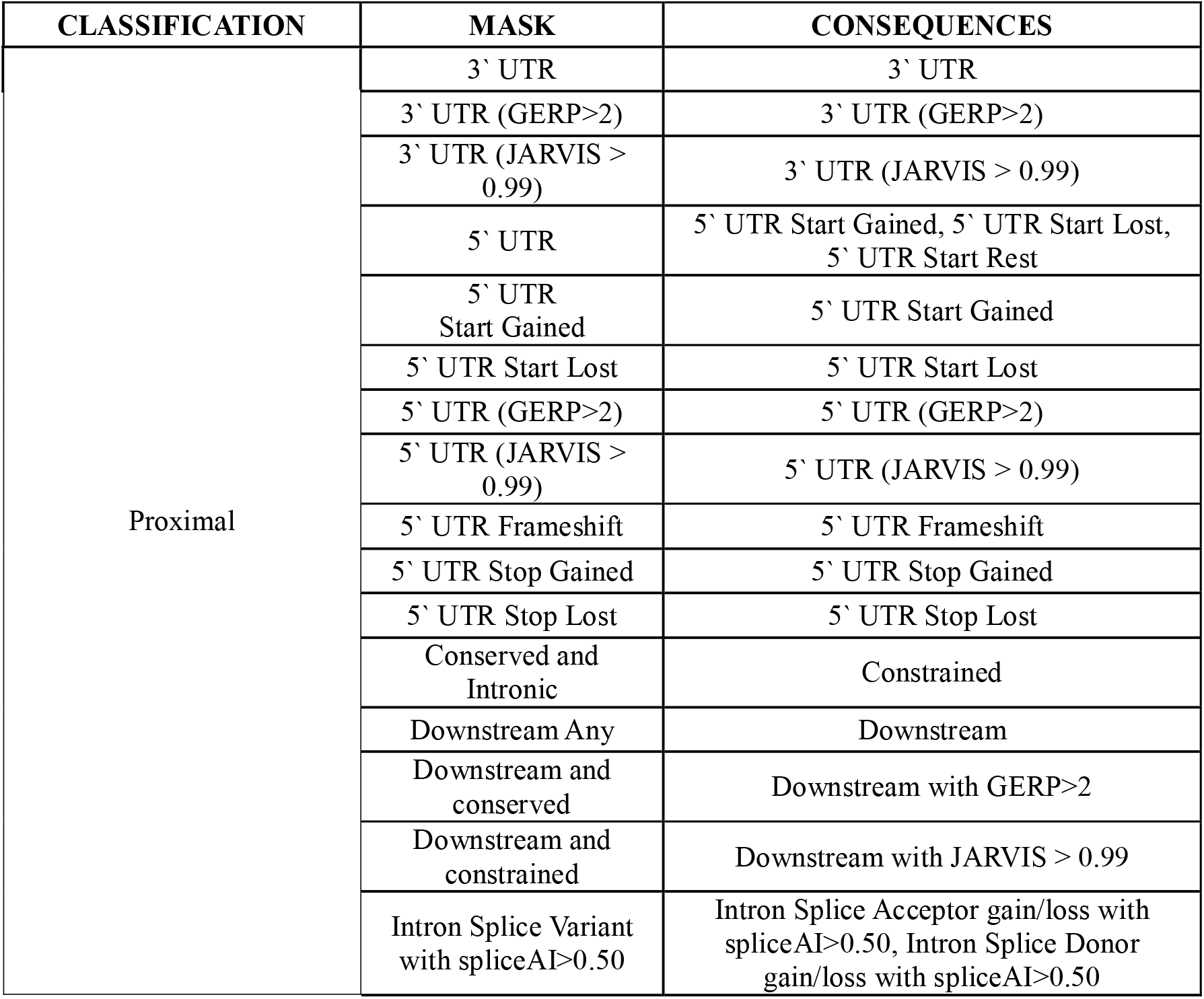
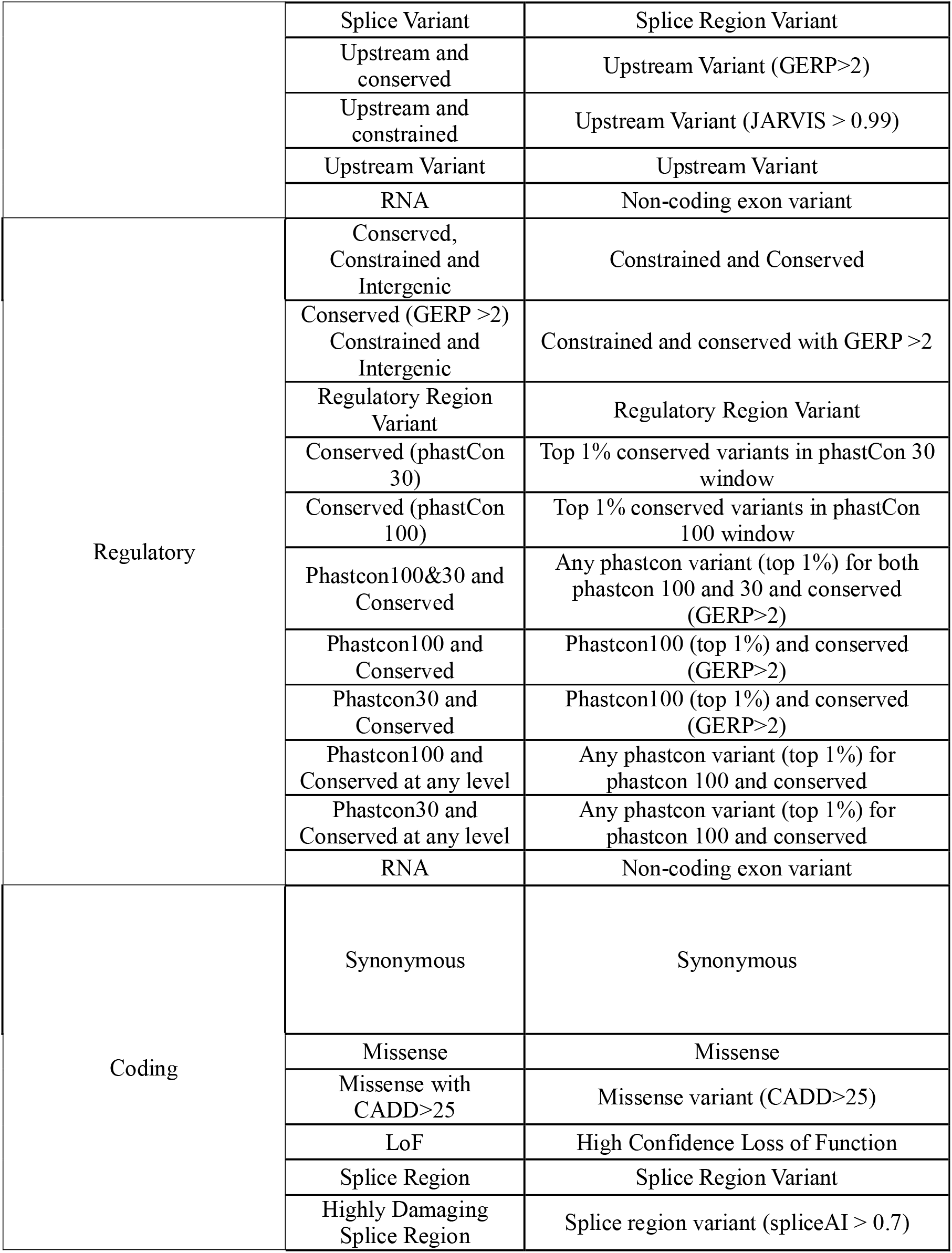

### Association Analyses

We performed both single variant and aggregate tests genome-wide for each of the three anthropometric phenotypes. All association analyses were corrected for age, sex, age squared, UK Biobank recruitment centre, the first forty genetic principal components and whole-genome sequencing batch.

### Single Variant Association Testing

To identify single variants associated with each phenotype we first performed an association test for all genetic variants with a minor-allele-count of at least 5 using *regenie*^*13*^ (v3.3) within 1,361 pseudo-linkage-independent chunks^40^ of the 22 autosomes, the two pseudo-autosomal (PAR; PAR1 & PAR2) regions of chromosome X and 100 equally sized chunks of the non-PAR regions of chromosome X. Chunk-wise lead variants were then selected in a conditional-joint analysis using an altered version of *GCTA-CoJo*^*14*^ (diff-freq = 0.2, cojo-p = 2.95×10^−19^), with the UK Biobank whole-genome sequencing data, limited to individuals with each phenotype, as an LD reference panel.

Testing revealed that *GCTA-CoJo* filters variants if their variance explained is >900-times the smallest variance explained by any independent variant (“sqrt(ldlt_B.vectorD().maxCoeff() / ldlt_B.vectorD().minCoeff()) > 30”: line 732 of gcta/meta/joint_meta.cpp at https://github.com/jianyangqt/gcta (accessed 17/03/2024)). We understand that this filter exists to capture statistical confounding caused by collinearity, for example if the reference genome used to calculate LD and the genetic data do not correlate well. However, for our purposes of jointly considering common and rare variants, where we have used the exact LD-reference panel matching our discovery data set, we found that this filter was falsely removing large-effect variants. We thus removed this filter and re-compiled *GCTA-CoJo*, which is available at (link to be added upon acceptance).

To define chromosome-wide independently associated variants, we applied a second round of the altered *GCTA-CoJo* algorithm, considering only those variants which were classified as independent at the chunk-level.

### Rare Variant Genomic Aggregate Testing

To identify coding and regulatory regions of the genome which were insufficiently powered for single variant analysis, we subsequently performed rare-variant (minor allele frequency <0.1%) genomic aggregate association tests using the annotations described in Table 1.

To test whether rare variant aggregate signals were caused by/confounded by residual LD and haplotype structure with common variants and or single variant signals we performed the following steps for each rare variant aggregate test result reaching *Bonferroni P* <0.05:

1. **To generate our primary discovery results** we adjusted for the common lead variants identified as independent signals in the joint (COJO) analysis (at MAF >0.1%)
2. **To identify independent aggregate associations**, if at least one aggregate passed our significance threshold we performed a forward stepwise regression. Starting from the most-strongly associated aggregate (by p-value), we performed an additional aggregate-testing run on aggregates reaching genome-wide significance, adjusting for all variants in the top signal. This process is repeated, with more variants added from the next most strongly associated aggregate, until no aggregate is genome-wide significant.

*TTN* was excluded from our conditional step-wise approach for aggregates due to regression convergence issues due to its size.

*regenie* performs four types of genome unit tests:

1. Standard BURDEN tests, under the assumption that each variant in a given gene unit mask has approximately the same effect size and sign on the phenotype
2. SKAT tests, where the sign of association of each variant in the unit is allowed to vary
3. ACAT tests, where the sign of association of each variant in the unit can differ, and only a small number of variants in the mask need be associated
4. ACAT-O, which is an omnibus test of BURDEN, SKAT and ACAT that aims to maximise the statistical power across the three tests

We performed each of the four statistical tests above for each mask for which a gene unit has at least one variant. Additionally, an association test was performed for all singleton variants (with MAC=1) in each unit. *regenie* also estimated an ‘all-mask’ association strength for each genome unit, which is an aggregation of the test statistics of the individual masks. To ensure that this did not result in a mixing of non-coding and coding association statistics, we split each gene transcript into a coding transcript, which we tested for all coding masks, and a proximal transcript that we tested for all proximal masks. Regulatory genome units were either classified by their ENSR assignment, by the extent of a 1kb constrained window, or a phastCon conserved window. We named sliding window masks by the region of the respective chromosome that they covered.

### Statistical Significance

Statistical significance was defined based on the minimum p-value observed for a whole-genome sequencing analysis of 20 randomly generated normally distributed continuous traits. The minimum p-value for single variant and aggregate association analyses were treated as independent: *P* (single variants) = 8.71 x10^−9^; *P* (aggregates) = 2.95 x10^−10^.

### Adjustment for previously reported GIANT loci

For each independent single variant and aggregate association, we performed a second round of adjustment for previously reported GIANT GWAS common-variant and exome-chip loci for each of the three anthropometric traits^2,19–23^. The effect sizes reported in the main results section are those calculated after adjustment for these previously reported loci – not those reported from the GCTA-CoJo analyses.

### Heritability

We calculated heritability using RHE-mc^41^ applied to UKB WGS data for N=343,451 maximally-unrelated European individuals, using variants with a minor-allele frequency greater than or equal to 1e-4. Maximally unrelated individuals were determined as the set of UKB-EUR-WGS for whom a GRM using all variants with MAF>0.1% had an off-diagonal component <0.05 (related individuals were removed in a way to maximise total sample size). For our primary analysis, we stratified the heritability into three minor-allele-frequency bins: 1×10^−4^ < MAF ≤ 1×10^−3^, 1×10^−3^ < MAF ≤ 1×10^−2^ and MAF>1×10^−2^). We additionally split variants into three bins of LD-score, calculated using GCTA^14^ (−-ld-score --ld-wind 1000 --ld-rsq-cutoff 0.01), LD-score≤ 25^th^ percentile, 25^th^ percentile < LD-score ≤ 75^th^ percentile; and LD-score > 75^th^ percentile. Where appropriate, we additionally split variants based on their inclusion within a 35kbp window surrounding known GWAS variants. In all cases, before running RHE-mc we rank-inverse-normalised each trait separately for each sex, and residualised on the following covariates: sex, age, age squared, centre, sequencing centre and genetic principal components 1-40. To minimise the risk of population stratification inflating our heritability estimates, we additionally adjusted for 100 haplotype components, calculated using SparsePainter^42,43^. As a secondary analysis, applied exclusively to height, we calculated heritability based on 17 non-overlapping minor-allele-frequency bins, from a minimum minor allele count of 10, with the following upper-bounds (inclusive): 1.7×10^−5^, 2.2×10^−5^, 3.7×10^−5^, 5.9×10^−5^, 1.0×10^−4^, 2.0×10^−4^, 4.0×10^−4^, 1.0×10^−3^, 5.0×10^−3^, 0.01, 0.02, 0.04, 0.1, 0.2, 0.3, 0.4, and 0.5.

