## Supplementary Information for "Whole-genome sequencing analysis of anthropometric traits in 672,976 individuals reveals convergence between rare and common genetic associations"

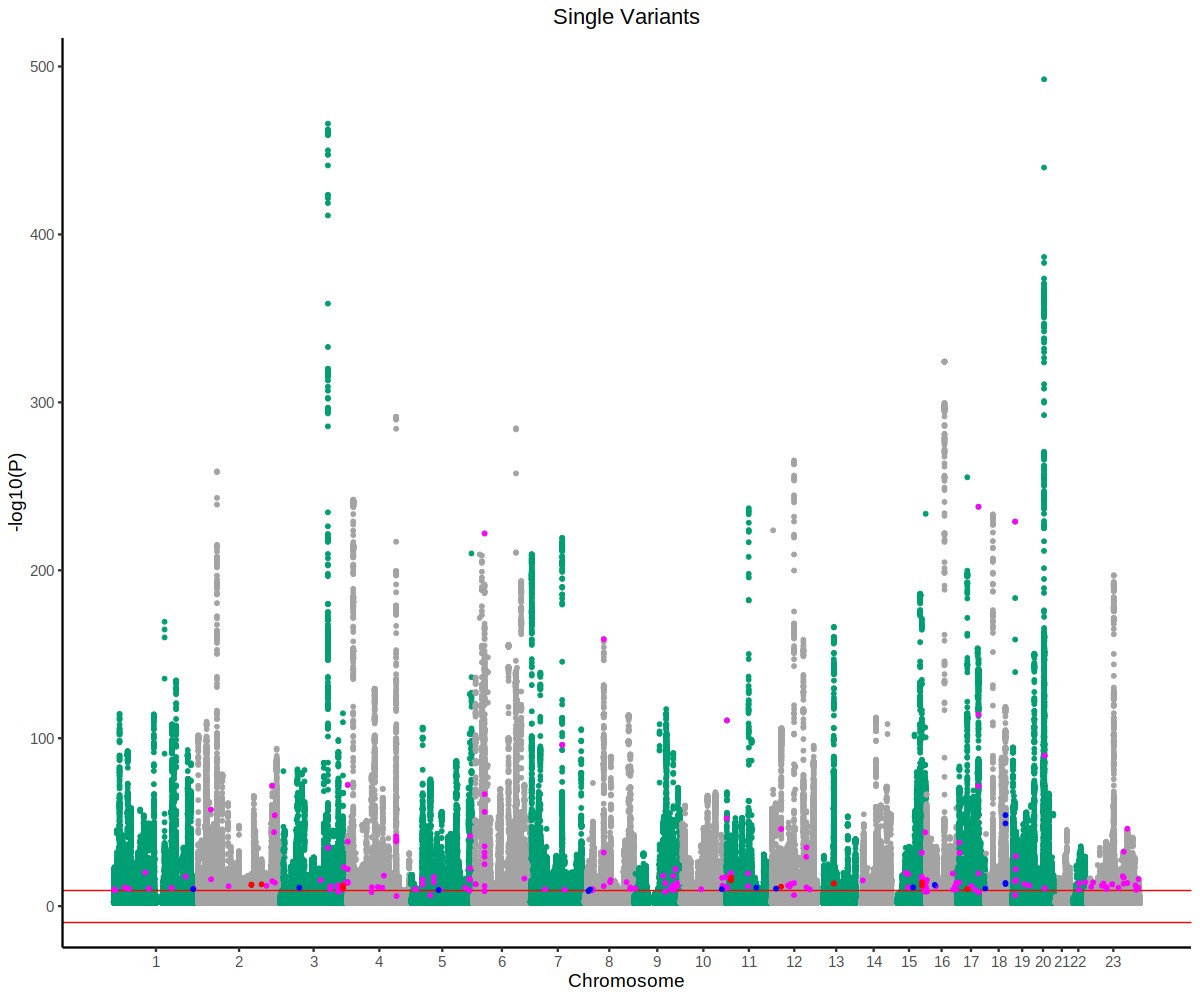

**Supplementary Figure 1: Manhattan plot of single variant associations with anthropometric traits in UKB.** Summary of all single variant associations for height, BMI and WHRadjBMI: the x-axis represents genomic position, segregated by chromosome and represented by alternating green and silver colours, on the x-axis, and -log10(P) on the y-axis. Jointly independent significant variants for each trait are uniquely coloured: magenta for height, red for WHRadjBMI and blue for BMI.

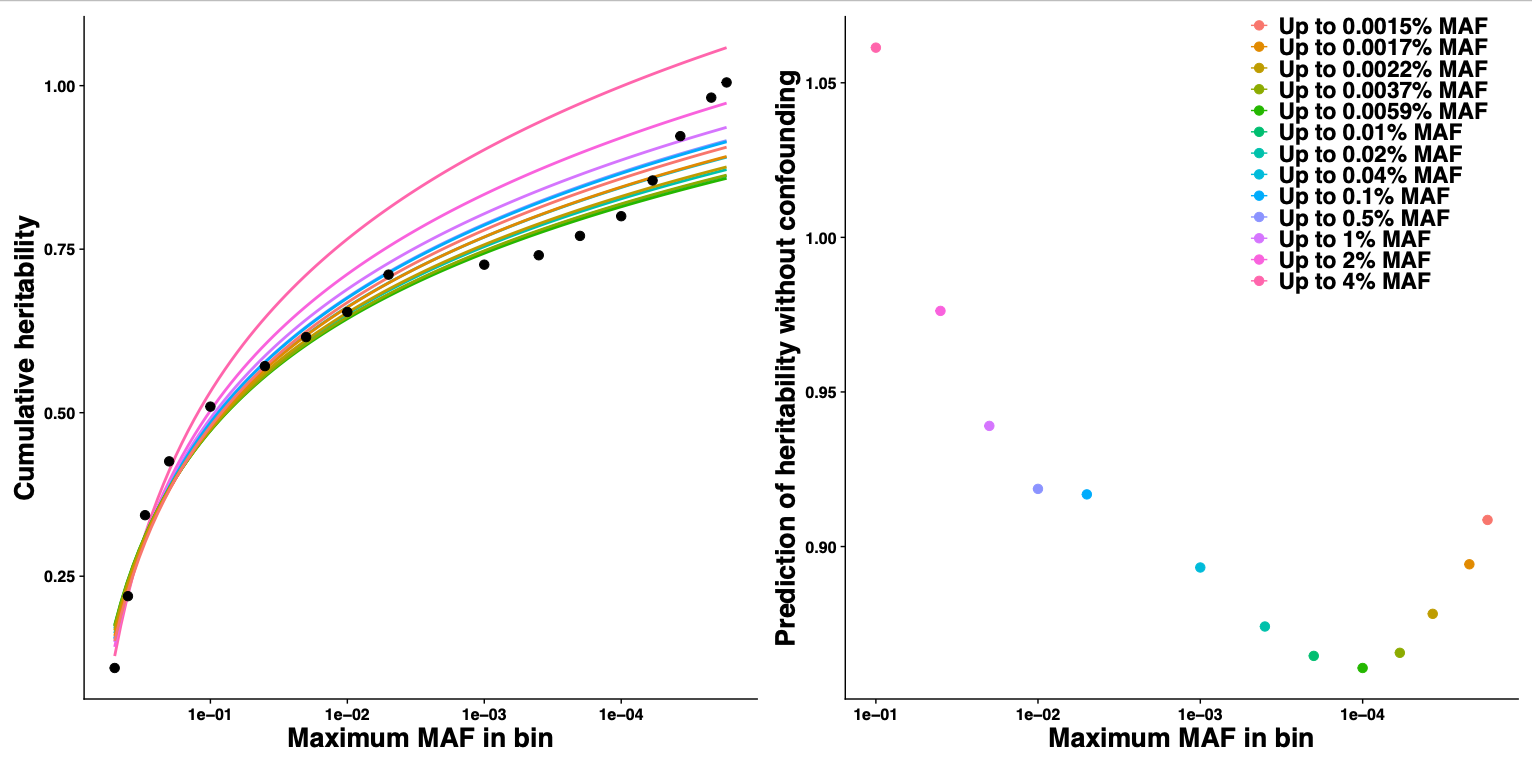

**Supplementary Figure 2: Estimating population stratification for heritability analysis** **Left:** Fitted logarithmic models (y = a*log(b*x)) for the relationship between minor-allele-frequency bin and heritability. Each model prediction (coloured line) is built from the inclusion of heritability estimates (black points) from increasingly lower MAFs (see legend). **Right**: estimation of total heritability predicted by logarithmic models (coloured by model)
